# Bacterial efflux pumps excrete SYTO^™^ dyes from bacteria and lead to false-negative staining results

**DOI:** 10.1101/2023.09.28.560001

**Authors:** G. A. Minero, P. B. Larsen, M. E. Hoppe, R. L. Meyer

## Abstract

Multidrug efflux pumps excrete a range of small molecules from bacterial cells. In this study, we show that bacterial efflux pumps have affinity for a range of SYTO™ dyes that are commonly used to label bacteria. Efflux pump activity will there lead to false negative results from bacterial staining and SYTO™ dyes should be used with caution on live samples.

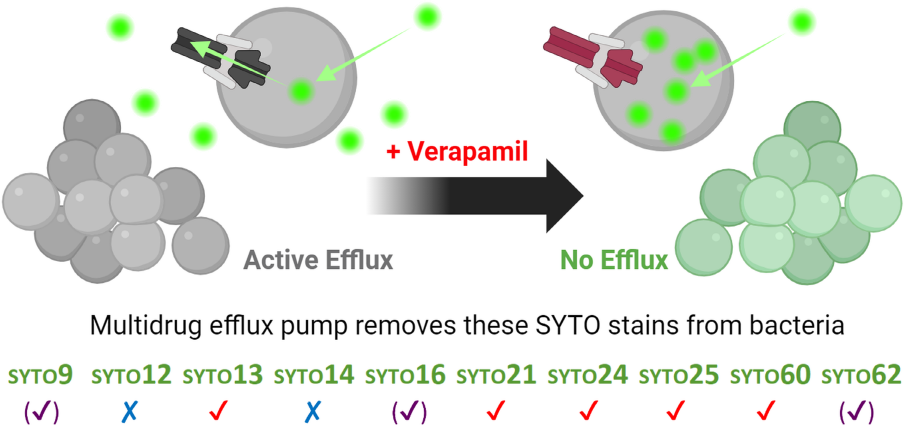

---

Efflux pumps (EPs) are mostly associated with transport of antibiotics and antimicrobials out of bacterial cells leading to efflux-mediated antibiotic resistance^1-3^ and can be induced in response to the chemical environment^2^. Drug-specific and multidrug EPs are common in bacteria and are either encoded on the chromosomal DNA or on a plasmid.

The activity of efflux pumps is often studied with fluorescence-based assays that detect the uptake and excretion of a fluorescent dye by an efflux pump with affinity for the specific dye. Such dyes include ethidium bromide (EtBr)^4^, Hoechst^5^, Nile red^5^, and SYTO™16 ^6^. Dye uptake occurs under conditions where the EP is inhibited, and the subsequent excretion is then measured after removal of the inhibitor^7^.

SYTO™ dyes are commonly used for fluorescence labelling of bacteria^8-10^, and since SYTO™16 was previously used to study multidrug EP activity in mammalian cells^6^, we hypothesized that bacterial efflux pumps also have affinity for SYTO™16 and other SYTO™ dyes. If this is the case, it will impact the result from SYTO™ staining of live bacteria. The aim of this study was thus to determine if bacterial multidrug efflux pumps can excrete SYTO™ dyes.

We use *Staphylococcus epidermidis* (*S. epidermidis*), a common culprit of implant-associated infections^11-14^, as a model organism to address this question, because it only encodes homologues of the S. *aureus* efflux pumps NorA/B/C^15^. The NorA efflux pump (388 aminoacid protein, 42 kDa) in *Staphylococcus aureus*^*7, 16, 17*^ and a NorA-like efflux pump in *Staphylococcus epidermidis*^*15*^ belong to the major facilitator superfamily and use the proton motive force to excrete fluoroquinolones and other antimicrobials from the cell^18^.

We used Verapamil (VRP) as the efflux pump inhibitor and EtBr as the fluorescent dye for detection of dye uptake and subsequent excretion by the efflux pump. Verapamil is an FDA-approved calcium channel blocker that potentiates the effect of several antituberculosis drugs through inhibition of the EPs *in vitro* and *in vivo*^19^. Verapamil is an effective inhibitor of NorA-EP^4, 20-22^ acting like an inhibitor of (i) Ca^2+^-channels and (ii) human P-glycoprotein^23^.

After confirming efflux pump activity with EtBr, we tested if the efflux pump also excreted SYTO™9, 13, 24, 25, 60, and 62.

First, *S. epidermidis* cells from overnight batch cultures (TSB media amended with 10 g/L glucose, 37°C, 180 rpm incubation, 3 biological replicates) were harvested by centrifugation (10.600 x g, 5 min) and re-suspended in mM9 buffer (Merck salts: 22 mM KH^2^PO4, 48 mM Na_2_HPO_4_, 8 mM NaCl, 19 mM NH_4_Cl, 0.4 mM MgCl_2_, 2 mM CaCl_2_) with 10 g/L Casa aminoacids (Thermofisher Scientific) and 10 g/L (approx. 50 mM) glucose^24^ (Merck) to OD_600_ = 1.5 and separated into 4 vials per biological replicate.

Next, 300 mg/L verapamil (V4629, VRP, Merck) was added to 3 vials, and 10 μM dye was subsequently added to all 4 vials.

The content of 2 vials (one with and one without VRP) was then transferred to a microwell plate (Nunc™ F96 MicroWell Black Polystyrene Plate), to quantify fluorescence in real-time (CLARIOstar BMG Labtech) as the dye was taken up by the cells (Figure 1A, left). If the NorA EP had affinity for the dye, the fluorescence would increase in the sample with VRP (where efflux is inhibited) compared to samples without VPR (where the dye is excreted).

**Figure 1.**
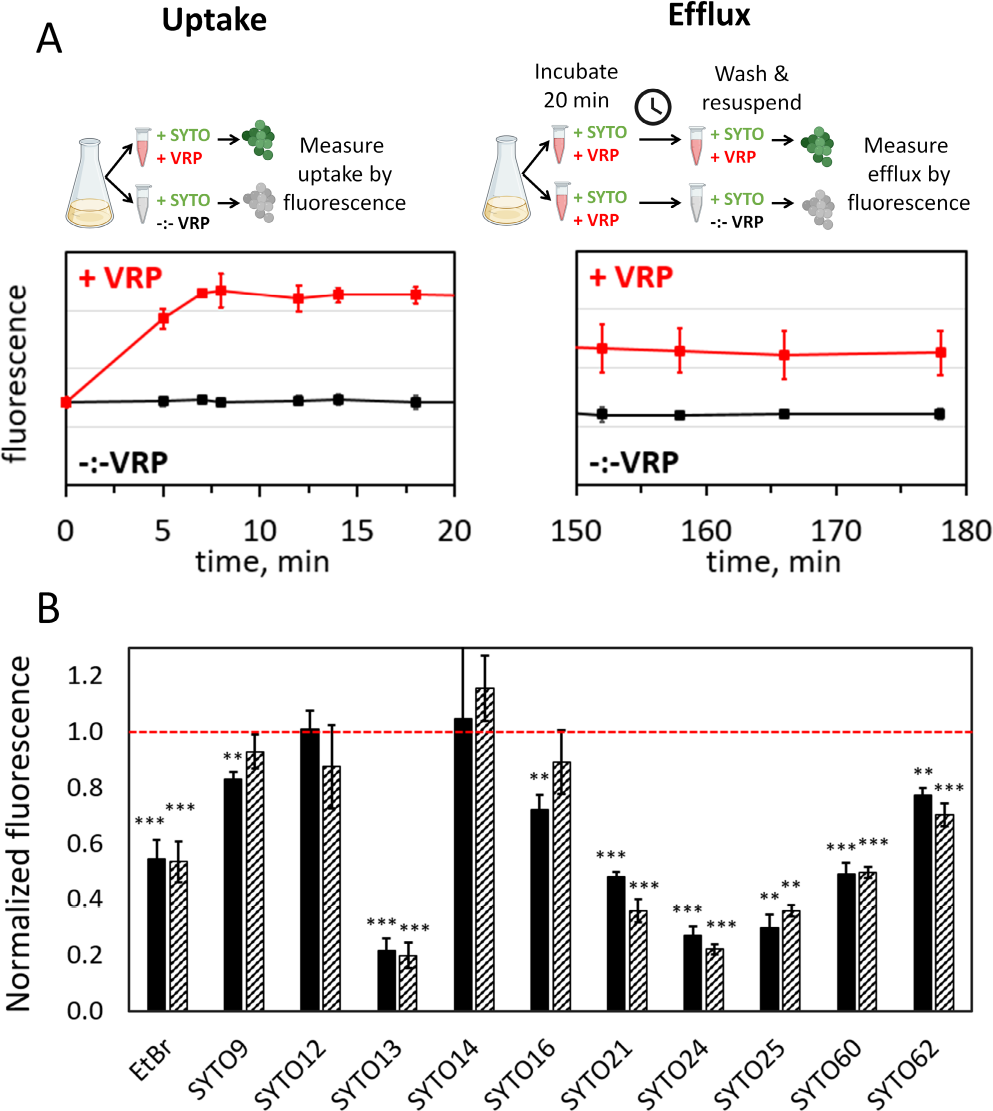
*S. epidermidis* multidrug efflux pumps have affinity for a range of SYTO™ dyes. A) Schematic illustration of the assay to quantify the effect of efflux pump activity on dye uptake and efflux, exemplified for SYTO^™^ 60. On the left, inhibition of efflux pumps by VRP results in dye uptake and increased fluorescence. On the right, both samples initially received VRP to facilitate dye uptake, but after a washing step, VRP was removed from one sample, resulting in dye efflux and decreased fluorescence compared to the sample with VRP. Identical fluorescence settings were used for detecting uptake and efflux. B) Mean fluorescence in samples with active efflux pumps relative to samples with VRP-inhibited efflux pumps (normalized pair-wise to the fluorescence in samples containing 300 mg/L VRP). Black bars reflect difference in dye uptake, and striped bars reflect difference in dye efflux. Values <1 signify that efflux pump activity impacts dye uptake (black bars) and dye efflux (striped bars) during uptake and washing, respectively. Student^’^ s t test, n=3, ^**^ =p<0.01.

The remaining 2 vials (with VRP) were incubated for at least 20 min to allow uptake of the dye before harvesting and washing the cells by centrifugation (10.600 x g, 5 min) and resuspension in mM9 buffer with 10 μM dye (one vial) or mM9 buffer with 10 μM dye and 300 mg/L VRP (one vial) followed by detection of fluorescence. Dye efflux was then measured as a decrease in fluorescence in samples without VRP compared to samples with VRP (Figure 1A, right).

Collectively, affinity of *S. epidermidis* efflux pumps for a given dye would be indicated by i) absence of dye uptake in samples without VRP, and ii) efflux of dye after transferring cells from a solution with VRP to a solution without VRP (Figure 1A, black curves). Figure 1A shows the data from an experiment with SYTO™60.

Figure 1B shows the fluorescence during uptake- and efflux experiments of samples with active efflux pumps relative to samples with VRP-inhibited efflux pumps. Values <1 signify that the EPs inhibit dye uptake (black bars) and facilitate dye efflux (striped bars). *S. epidermidis* efflux pumps had affinity for SYTO™13, 21, 24, 25, and 60. It had some affinity for SYTO™9, 16, and 62 resulting in slightly impaired dye uptake, while there was no effect on SYTO™12 and 14.

In addition to bulk fluorescence measurements, we also visualized the fluorescence from individual cells using confocal laser scanning microscopy (CLSM, Figure 2). This analysis was performed on samples incubated with EtBr and SYTO™ 9, 13, 24, 25, and 60. The four images show uptake and efflux of dyes and were captured with the same acquisition settings such that fluorescence intensity could be compared among samples with the same dye. CLSM imaging corroborated the conclusion from bulk fluorescence, namely that EP activity affected bacterial staining by most of the tested SYTO™ stains, which strongly affects detection of the cells by microscopy. Quantification of single-cell fluorescence intensity showed that fluorescence from bacteria with active EPs compared to inhibited EPs was 0.24 ± 0.01 for EtBr (n=3) and 0.16 ± 0.05 for SYTO™60 (n=3), which were both effectively excreted. The value was 0.71 ± 0.22 for SYTO™9 (n=3), showing a smaller yet significant reduction in fluorescence intensity caused by the activity of efflux pumps. SYTO™9 is commonly used in Live/Dead staining of bacteria. Efflux of SYTO™9 will not lead to false identification of dead bacteria in a sample, but it could lead to under-representation of the amount of live cells.

**Figure 2.**
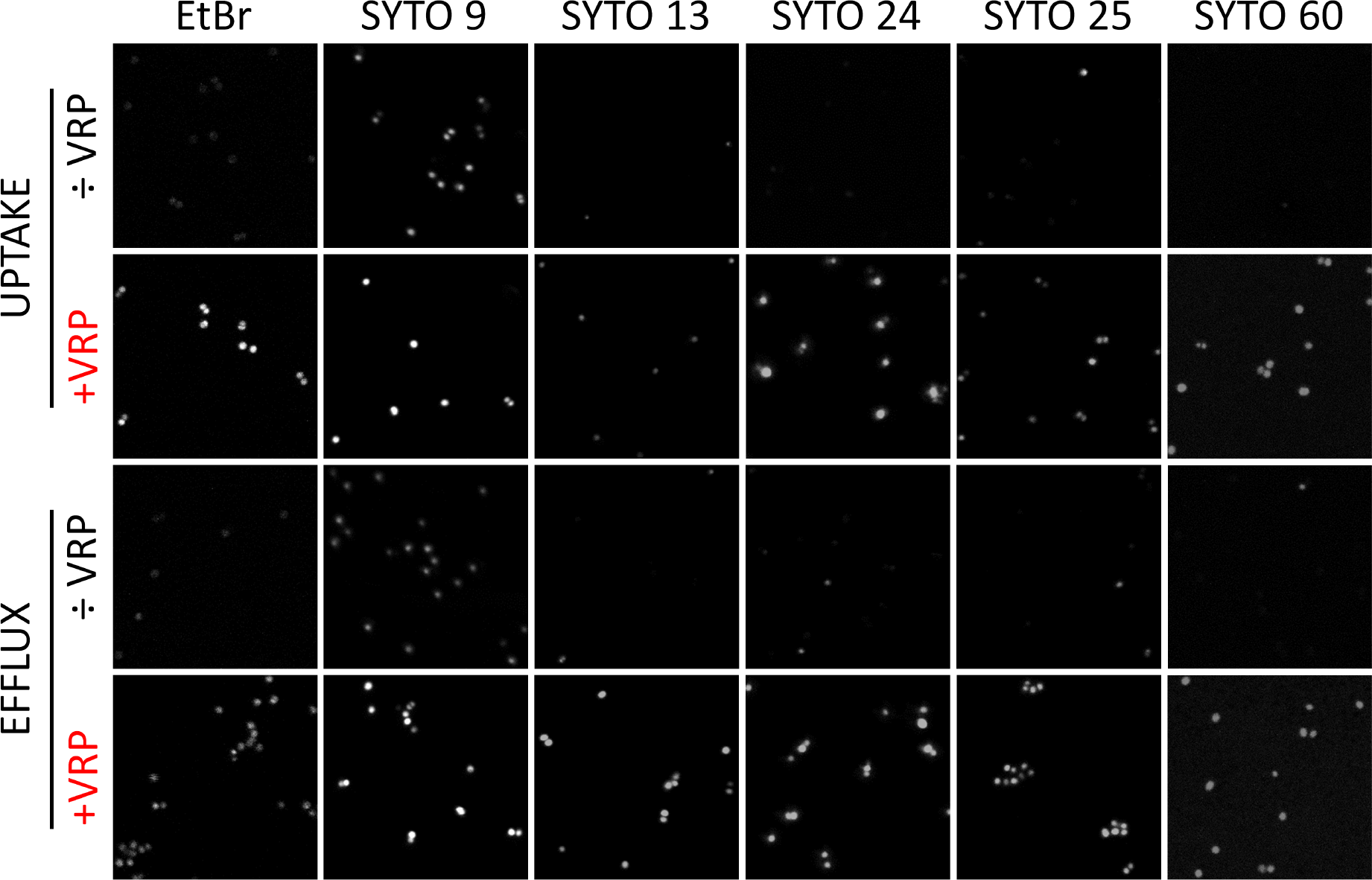
Glucose (10 g/L) fuels Efflux pump in *S. epidermidis* planktonic culture while verapamil (300 mg/L) inhibits the Efflux pump activity. 2D CLSM images (25.3 x 25.3 μm) of selected group of samples (see Figure 1 for the details of sample treatment) containing 10 μM EtBr, SYTO^™^ 9, SYTO^™^ 13, SYTO^™^ 24, SYTO™ 25, and SYTO^™^ 60.

In conclusion, we show for the first time that a wide range of SYTO™ dyes are excreted from *S. epidermidis* cells when efflux pumps are active. This means that such SYTO™ stains can be used to detect efflux pump activity, but importantly, it shows how efflux pump activity affects cell staining for use in e.g. microscopy or flow cytometry. SYTO™ staining should be used with care, and we think that efflux pump inhibitors can improve sensitivity and reproducibility of SYTO™ staining in unfixed samples.

We thank the Villum Foundation (Grants no. 00028321 and no. 00050284) for funding of this work. We also thank Dr. Ebbe Sloth Andersen for providing us with an access to the fluorescence measurements at the CLARIOstar plate reader.

## Author Contributions

G. A. Minero performed supervision of the experimental work, data analysis and presentation, as well as drafting and revising this manuscript. P. B. Larsen and M. E. Hoppe performed the experiments and revised the manuscript. R. L. Meyer performed conceptualization as well as supervision of this work and revision of the manuscript.

